# Previous motor task performance impacts phase-based EEG resting-state connectivity states

**DOI:** 10.1101/2023.07.06.547927

**Authors:** Nils Rosjat, Maximilian Hommelsen, Gereon R. Fink, Silvia Daun

## Abstract

The resting human brain cycles through distinct states that can be analyzed using microstate analysis and electroencephalography (EEG) data. This approach classifies, multichannel EEG data into spontaneously interchanging microstates based on topographic features. These microstates may be valuable biomarkers in neurodegenerative diseases since they reflect the resting brain’s state. However, microstates do not provide information about the active neural networks during the resting-state.

This article presents an alternative and complementary method for analyzing resting-state EEG data and demonstrates its reproducibility and reliability. This method considers cerebral connectivity states defined by phase synchronization and measured using the corrected imaginary phase-locking value (ciPLV) based on source-reconstructed EEG recordings. We analyzed resting-state EEG data from young, healthy participants acquired on five consecutive days before and after a motor task. We show that our data reproduce microstates previously reported. Further, we reveal four stable topographic patterns over the multiple recording sessions in the source connectivity space. While the classical microstates were unaffected by a preceding motor task, the connectivity states were altered, reflecting the suppression of frontal activity in the post-movement resting-state.

## Introduction

The state of the human brain at rest has been studied extensively using multichannel electroencephalography (EEG) and functional magnetic resonance imaging (fMRI), showing that the brain stays active in an organized manner for unpredictable incoming stimuli (1–3). Connectivity analyses identified several networks, including the visual, sensory-motor, basal ganglia, and default mode networks (4–7). In fMRI analyses, these networks are extracted by analyzing the correlations of BOLD signal fluctuations among various brain regions (4). However, the fluctuations found in these signals are too slow to be associated with preparations or responses to unpredictable stimuli.

On the other hand, analysis of EEG signals providing a much higher temporal resolution focuses on the correlation of fluctuations in amplitudes of oscillatory activity (8; 9). Several methods extract EEG signal information to analyze resting-state networks. For example, the so-called microstate analysis is a well-established method that examines the topographical configuration of electrode potentials as measured at the scalp level. Recent advancements in the study of EEG microstates have expanded our understanding of their number and characteristics. While traditionally, four prototypical microstates have been consistently identified (e.g. (10–12)), emerging evidence suggests a more nuanced picture. Specifically, a comprehensive meta-analysis by (13) and a systematic review by (14) have highlighted the potential existence of up to 6 or 7 reliably identified microstates. These microstates spontaneously transition between each other within several hundreds of milliseconds. This rapid transition most likely reflects the switching between neural networks. For instance, pre-stimulus microstates have been shown to affect stimulus response performance, suggesting that EEG signals unravel the role of brain microstates when preparing for incoming stimuli (15). Although combined EEG-fMRI studies indicate that the EEG microstates are highly correlated with fMRI resting-state networks (11), the neural mechanisms underlying microstate activity, their functional relevance, and the dynamic and complex nature of the rapid transitions remain elusive.

In this study we took an approach different from the microstate analysis to investigate the neural mechanisms underlying active brain networks at rest. For this, we aimed to define and analyze connectivity states underlying resting-state networks using EEG resting-state data. We furthermore tested whether these networks are stable over several measurement days and whether they are affected by cognitive or motor tasks. Thus, we investigated whether the acquisition time within an experiment, i.e., pre- or post-task performance, affected the resting-state connectivity states. We used source reconstruction methods to determine the source activity at any given time and determined the sources’ connectivity state using the corrected imaginary phase-locking values (ciPLV). This approach is to be distinguished from work by (16). In their study, the authors utilized microstates to parcellate the resting-state time course into brain states before conducting connectivity analysis on the microstates. In contrast, we compute resting-state connectivity in source space and cluster the obtained source networks into distinct groups, which we refer to as connectivity states. This approach enables us to examine the topographical organization of the source networks. It addresses previous concerns regarding the combination of microstate analysis and connectivity analysis, pointed out by (17). Thus, while previous literature aimed to inform connectivity analysis by microstate segmentation, we used a fundamentally different approach and strive to provide a novel perspective on understanding EEG resting-state connectivity. We thus extend previous work by offering an alternative approach that enhances our understanding of the neural dynamics underlying resting-state brain activity.

Many studies have identified disruptions in specific microstate features (e.g., occurrences, coverages) in neuropsychiatric disorders, including schizophrenia (18; 19), dementia (20), depression (21), Tourette’s syndrome (22), or panic disorder (23). In contrast, data on stroke-related disruptions of microstates or active resting-state networks remain sparse. The source-localized connectivity-state approach presented here might help to better understand disturbed cortical network organization following a stroke. Before applying this method to patients, it is, however, mandatory to test whether the results in healthy subjects are stable across measurements and time.

The methodological validation emphasized in this study ensures that subsequent research can rely on these methods as being robust. In clinical settings, especially in studies involving stroke patients or other neurological disorders, this research can pave the way for more personalized treatment approaches by analyzing single subject connectivity patterns and their deviations from healthy connectivity states. By understanding individual connectivity patterns, future studies can tailor interventions to patients’ specific needs, potentially improving recovery outcomes.

## Materials and Methods

### Participants

In this study, 27 healthy young subjects (11 female, age (mean ± sd): 27.9 years ± 3.4, range: 22-34 years) participated and received monetary compensation. Participants had normal or corrected-to-normal vision, no neurological or psychiatric disease history, were not under medication at any time, and had no cranial metallic implants. According to the Edinburgh Handedness Inventory (24), 22 participants were right-handed (score > 50), two were left-handed (score *< −*50), and three had intermediate scores. The study was approved by the Ethics Commission of the Faculty of Medicine, University of Cologne (ID: 14-006). All participants provided written informed consent before the start of the experiment.

After applying all exclusion criteria (see below), datasets from 24 (of the 27) participants were used for statistical analyses.

### Experimental Setup

The software Presentation (v. 20.2 Build 07.25.18, Neurobehavioral Systems, Inc.) was used for the visual representation of the experimental paradigm. The behavioral responses of the subjects were recorded via an fMRI-compatible button pad (1-hand) system (LXPAD-1×5-10M, NAtA Technologies, Canada).

During the experiment, a 64-channel scalp EEG was recorded with active Ag/AgCl electrodes (actiCap, Brain Products GmbH, Germany). The electrodes were positioned in the standard 10-20 layout (ground AFz, reference on left mastoid). Three electrodes (FT9, FT10, TP10) were used for EOG recording. One electrode was placed under the left eye for vertical movements and one each near the left and right lateral canthi for horizontal movements. The recorded voltages were amplified during the measurements by a BrainAmp DC amplifier (Brain Products GmbH, Germany).

A stereotactic neuronavigation system (Brainsight v. 2.3, Rogue Research Inc., Canada) was used (for details, see (25)) to ensure reliable positioning of the EEG electrodes over the multiple measurements taken on different days.

The experimental sessions took place in a controlled environment to ensure minimal external influences on the participants’ resting-state. The chamber used for the experiments was dark, sound-proof, and magnetically shielded. This setup aimed to minimize visual, auditory, and electromagnetic interferences that could potentially impact the resting-state measurements. During the experimental sessions, participants were seated upright, facing a computer screen.

### Experimental Protocol and Paradigm

Subjects were each measured at identical times of the day for five consecutive days (Monday through Friday). For one subject, the fifth measurement had to be performed three days apart for technical reasons. For all subjects, each session (i.e., each measurement day) consisted of two resting-state measurements interrupted by a non-rest finger-tapping task. Besides, a more complicated finger movement sequence was recorded at the end of the session, which will not be considered further here.

The individual conditions were organized in detail as follows and together lasted for about 40 minutes.

1. Resting State (RS1): A white dot was shown in the center of the screen for the entire duration of 5 minutes. Subjects were instructed to keep their eyes open, fixating their gaze on the white dot. Further, it was emphasized that the subjects should relax, avoid movements as much as possible, and not think of anything specific. This task will be referred to as RS1.
2. Tapping Task (Tap): During this task, a white dot was also presented for fixation on the center of the screen. However, the dot faded in this task every 10-14 seconds. During this period, subjects had to press the button on the response pad with their left index finger as quickly and as often as possible until the dot reappeared on the screen. The entire block consisted of 60 movement periods (dot absent, Move) and 60 waiting periods (dot present, Wait) and lasted in total about 14 minutes.
3. Resting State (RS2): The second block of the Resting State (RS2) proceeded as the first.
4. Sequence Task (Sequence): In this task, the subjects had to press a given, more complex sequence as quickly and as often as possible with the fingers of one hand. This task will not be described in more detail here, as it will not be considered in the following analyses (see (25) for details).

### EEG Preprocessing

EEG preprocessing was performed using EEGLAB (26), integrated into MATLAB (R2016b, Math-work Inc, Natick, MA). EEG source reconstruction was carried out using MNE-Python (v. 0.22) in Python (v. 3.8.8). Further analyses were performed using custom scripts also implemented in Python.

The continuously recorded data was first downsampled to a sampling frequency of 128 Hz. In the next step, the data were band-pass filtered to 1-40 Hz frequency with a Hamming windowed sinc FIR filter (i.e., high pass followed by low pass). The continuous recordings then underwent an artifact correction process to remove oculomotor activity related to eye blinks and saccades based on the information from the EOG electrodes. Eyeblink removal was performed for each day separately on a copy of the data set following the procedure introduced by (27). ICA weights obtained from this procedure were transferred and applied to the original dataset. Independent components related to eyeblinks and saccades were identified using the automatic detection algorithm ADJUST (28), implemented as a plug-in for EEGLAB.

Following eyeblink correction, the original dataset underwent visual inspection to remove time-intervals and channels contaminated with artifacts. Signals of rejected channels were interpolated by using information from other channels using spherical spline interpolation. Time-intervals showing artificial behavior were cut out entirely from the data. All channels were then re-referenced to the Common Average Reference (29).

### Data Quality Assessment

In order to ensure data quality and minimize the potential influence of drowsiness on the resting state measurements, we carefully checked for any signs of decreased vigilance or wakefulness in the subjects. If epochs of drowsiness were identified, they were excluded from further analysis. This approach aimed to maintain the validity and reliability of the resting state measurements and minimize any confounding effects arising from decreased vigilance or wakefulness.

After preprocessing, we obtained 135 data sets (27 subjects x 5 days). All subjects completed at least the first three tasks, i.e., RS1, Tap, and RS2 (see (25)). To be included in the final data analysis, at least 4 of the 5 subjects’ data sets had to meet the following quality criteria:

1. Fewer than 7 excluded channels,
2. *>* 180 s artifact-free RS1,
3. *>* 180 s artifact-free RS2.

Of all 27 subjects, 24 met the above criteria and were included in the analyses reported below. Of these, 18 (of 24) had performed all 4 tasks, and the remaining 6 (of 24) had performed the first 3. The best 4 of the 5 sessions (days) were used from all subjects to ensure a comparable amount of data per subject. For subjects with 5 good sessions, the first day of each measurement was excluded.

### Microstate Analysis

We determined classical microstates at the electrode level for face validity before analyzing the source activity. For this purpose, we used the MNE Microstates Toolbox (30). The continuous recordings at the electrode level were divided into 4 states by applying the k-means algorithm with *k* = 4, each with 10 different initializations per subject and measurement. The best of the 10 fits for each subject was used to determine the 4 common group centroids.

While recent studies have proposed the existence of more than four microstates, our choice of constraining our microstate clustering to four states was based on specific observations from our dataset. During our analyses, we noticed that as we increased the number of states beyond four, several resultant states exhibited high similarity and were not distinctly separable. The four-state solution balanced between capturing the essential dynamics of the data and ensuring the clear interpretability of our results.

### Source Localization

The continuous recordings were projected from the electrode onto the source space using dynamic statistical parametric mapping (dSPM) (31). Afterward, the norm of the source orientations was taken (32). We applied methods from FreeSurfer for source-space reconstructions (33; 34). As no individual MRIs were available for the subjects, source data were mapped onto FreeSurfer’s “fsaverage“ common template source space. Source locations were then grouped into 150 parcels according to the Destrieux Atlas (35). For each parcel, one common, i.e., shared, parcel time course was extracted by finding the dominant direction of the source space normal vector orientations within each parcel. Each time series at vertices whose orientation is more than 180° different from the major direction underwent a sign-flip. Finally, all vertices within each parcel were averaged to produce the final common source time course. Source localization was performed on the movement data first to ensure the proper functioning of the algorithm, as the sources are known in this condition.

### Network Construction

Following source localization, the procedure for creating the network was divided into two steps: (a) connectivity analysis, and (b) graph construction (see Fig. 2).

**Fig 1.**
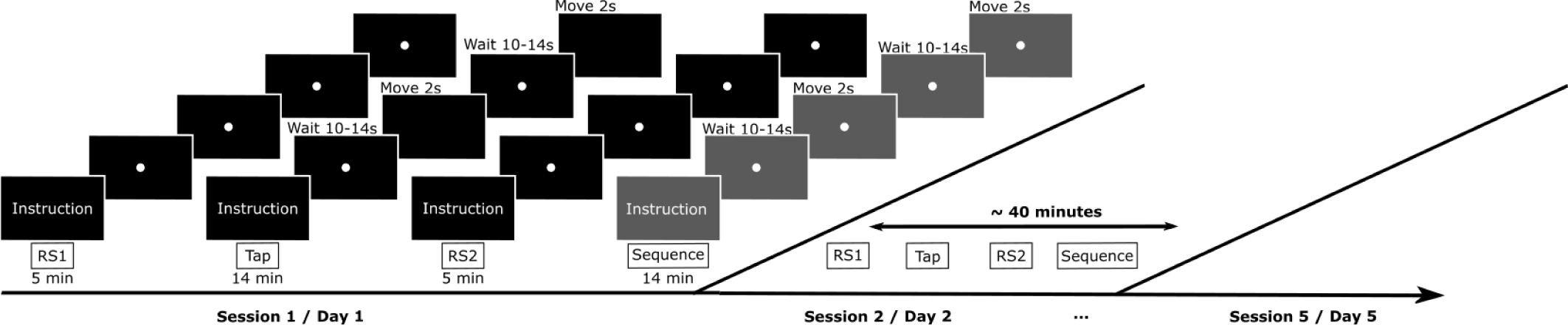
Experimental Paradigm. Figure showing the experimental conditions. Top: Single measurement sequence consisting of 5 min Resting State (RS1), 14 min Tapping Task (Tap), 5 min Resting State (RS2), and, finally, 14 min Sequence Task (Sequence). The sequence task is shown in grey as it will not be considered for further analyses; Bottom: Temporal sequence of measurements for day 1 to day 5.

**Fig 2.**
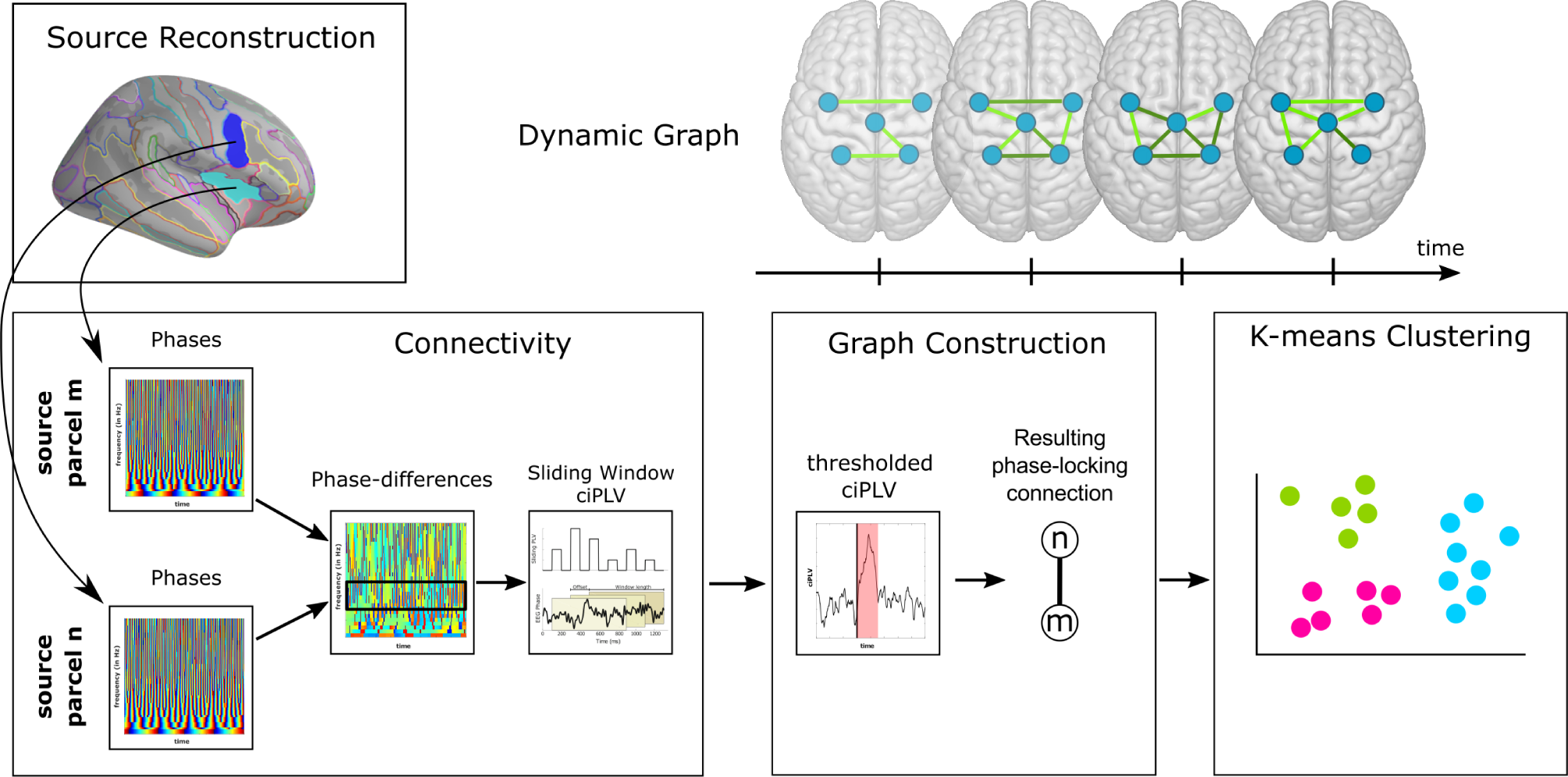
Connectivity State Pipeline. Representation of how the connectivity states were determined. Top row (left): extraction of source activity for two parcels, m and n; bottom row (left): computation of the ciPLV connectivity metric; bottom row (middle): graph construction for each time window by thresholding the ciPLV; top row (right): depiction of the resulting dynamic graph networks; bottom row (right): application of k-means clustering for connectivity state assignment.

(a) **Connectivity Analysis:** In this step, each parcel’s phase was extracted and a phase-synchronization analysis applied. The objective was to quantify the connectivity strength between all possible pairs of parcels throughout the experiment’s duration.

(b) **Graph Construction:** Upon establishing the connectivity strengths, this subsequent step employed thresholding techniques to determine the strongest connections at each specific timepoint. The graph construction was performed for each timepoint to represent the dynamic changes in source interactions over the whole time.

These procedures are detailed below, providing a comprehensive overview of the methodologies employed.

First, we transformed the continuous source time courses of the resting-state data to the time-frequency domain using Morlet wavelets (36). We focused our analysis on the alpha frequency range (8-12 Hz) and applied a step size of 0.5 Hz. The choice of the alpha frequency band was based on several reasons outlined below:

1. Importance for motor networks: The alpha band is relevant (37; 38). By focusing on the alpha band, we aimed to explore potential connectivity patterns and network dynamics specific to motor-related brain regions.
2. Connectivity analysis: It is necessary to narrow the frequency range of interest for phase connectivity analysis. Broadband approaches may not be suitable for capturing specific connectivity patterns effectively. By focusing on the alpha band, we aimed to target a specific frequency range that is known to exhibit meaningful connectivity patterns.
3. Microstate analysis: Previous studies utilizing classical microstate analysis have reported consistent states for broader frequency ranges and alpha filtered data during resting-state measurements (e.g., (39)).
4. Technical constraints: In our study, we faced technical constraints that influenced our choice of frequency boundaries. Lowering the frequency boundary would have resulted in decreased time resolution, which would have hindered our ability to capture fine-grained temporal dynamics and connectivity changes.

By narrowing our focus to the alpha band, we aimed to balance between the methodological requirements of the connectivity analysis, the existing literature on alpha oscillations, the relevance of the alpha band to motor networks, and technical considerations associated with our data processing pipeline.

The Morlet wavelet cycles were set to increase from 4 at the lower-frequency boundary to 6 at the upper-frequency boundary. After the time-frequency transformation, we extracted the time courses of the amplitudes and phases for each frequency. We quantified the connectivity between two different parcels (regions) by synchronization determined by the single-trial corrected imaginary phase-locking value (ciPLV; (40)). As we used continuous data without any stimulus onset or reoccurring trials, we followed the suggestion by Lachaux et al. (41) and computed the phase-locking over a specific period instead of trial-wise phase-similarity. Thus, two parcels are defined as being connected if the phase difference remains constant over this period. In our analysis, this period was set to 300 ms. For further details on the choice of the length of this period, please refer to the Supplementary Materials, Fig. S1.

For a pair of parcels m and n, ciPLV is defined as:

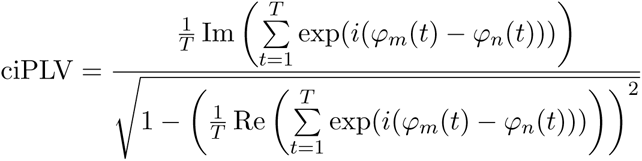

Here *φ_m_*(t) denotes the phase of the source reconstructed EEG signal at parcel m at time point t. T is the total number of time points in the 300 ms interval. The ciPLV is computed in a sliding window approach to obtain a time-dependent connectivity measure. Previous literature (e.g., (39)) has shown microstates to be stable for predominantly 100 ms before transitioning to the next state. Thus, the windows were set to have a 200 ms overlap to achieve a time resolution of 100 ms. While a minimum ciPLV = 0 represents a random distribution of phase difference over the whole time interval, a maximal ciPLV = 1 occurs only in the case of perfect phase-locking of the phase differences with a phase difference of *π*/2. Connectivity with zero-phase lag is intentionally removed from the analysis by the specific selection of ciPLV as the connectivity measure since it is, to a high degree, artificially created by source leakage.

The connectivity analysis classifies pairs of parcels at each time window into those with higher and lower degrees of synchronization. In the following, we will focus on the 10% most robust connections at each given time window with a resolution of 100 ms. By focusing on the top 10 % of connections, we emphasize the most robust and consistent interactions, which helps reduce noise and the potential influence of spurious links. Such a sparse representation is often preferred in network analyses to enhance the network’s structure clarity and emphasize the most salient features. Following these steps, we can define a binary undirected graph *G_t_* = (*V, E_t_*), where the graph G is defined by a set of vertices V, i.e., the source locations, and edges *E_t_* : *V × V → R*, i.e., the strongest connections at each time window. The dynamic graph, also called a dynamic network, is then defined as an ordered set of graphs *G*(*t*) = *{G_t_|t ∈* [1*, · · ·, T_tot_*]*}*. Here, *T_tot_* denotes the total amount of artifact-free resting state data for a given participant. The use of dynamic graphs is crucial here, as they account for time-varying connectivity patterns necessary to determine the time-evolving formation of subnetworks that a static network could not unravel.

### Connectivity State Clustering

For clustering the connectivity patterns into different connectivity states (CS), we used a similar approach as described above for the microstate analysis. For one, we determined 4 different connectivity states by applying the k-means clustering algorithm on the dataset, consisting of 24 subjects and 4 measurement days for RS1 and RS2. Additionally, we used the elbow method (42; 43) to determine the optimal number k for the k-means algorithm. This approach led to 8 connectivity states, which we computed for all subjects (24), measurement days (1–4), and tasks (RS1, RS2). In our analyses, we explored both 4-state and 8-state solutions for connectivity patterns, reflecting our interest in capturing the granularity regularly reported in the resting-state EEG literature (e.g., (10–12; 44)). While the 8-state solution provided additional states, we observed several similarities in connectivity patterns. Given these observations and the precedence in the literature, we primarily focused on the more distinct 4-state solution for clarity in our findings. Details on the results for the 8-state solution are provided in the Supplementary Material.

### Statistical Analysis

The coverages of the microstates were compared using pairwise t-tests. Since the distributions of the connectivity state coverages did not fulfill the requirements of normally distributed data those states were compared using the Wilcoxon signed-rank test within each RS measurement. This non-parametric test assesses whether two related paired samples originate from the same distribution. Specifically, it examines whether the distribution of differences *x − y* is symmetrically centered around zero.

We performed additional Wilcoxon signed-rank tests for each connectivity state to further compare the effects of measurement days (1–4) or pre- and post-motor performance resting-state (RS1 vs. RS2).

Besides the Wilcoxon signed-rank test, we applied the Intraclass Correlation Coefficient (ICC) to evaluate the reliability of our metrics across multiple measurement days. The ICC assesses the consistency and agreement of measurements taken on different occasions. A higher ICC value indicates greater reliability of the measurements.

These statistical analyses provide insights into the differences and consistencies between various measurements and conditions in our study.

## Results

### Microstate Analysis

In the first step, we performed a classical microstate analysis of the resting-state data (RS1 and RS2) using *k* = 4 clusters. We added this step to investigate the variability of the microstates over multiple measurement sessions on different days. Further, we tested whether the performance of a motor task before a resting-state influences the microstates. We conducted the microstate analysis in two frequency ranges: (i) a focused alpha frequency range of 8-12 Hz, analogous to our connectivity state analysis, and (ii) a more conventionally used broad range of 1-40 Hz, prevalent in the microstate literature (14). Given the strong agreement in the outcomes from both approaches, we will subsequently only present the results obtained from the 1-40 Hz frequency range for clarity and continuity of the established standards.

In Fig. 3A, we show the four microstates determined from our dataset. The four states exhibit the same basic topographies as have been reported by many studies in the previous literature (10–12). The four microstates found in our data are characterized topographically by (i) a diagonal polarity shift from left parietal to right frontal regions (MS A), (ii) a mirrored diagonal polarity shift from the right parietal to left frontal regions (MS B), (iii) a polarity shift at the central electrodes (MS C), and an occipital polarity shift (MS D). When separating the sessions into pre-task (RS1) and post-task (RS2) resting states, there were no apparent changes in the four microstates (see Supplementary Material, Fig. S2). Similarly, the microstates remained stable when comparing the different measurement days (session 1 to session 4; see Fig. S2). The analysis of the coverages of the four microstates showed no significant difference between all four microstates (Fig. 3B). When looking at the transition between the four microstates, we can see that each state had a high probability (*>* 50 %) of staying in the same state while transitions to the other states were evenly distributed at roughly 15 % (Fig. 3C). Thus, there existed no preferred microstate at the scalp level and no clear order of state transitions. This result is stable across the 4 measurement days (Fig. 3D) where no significant difference was found between the days.

**Fig 3.**
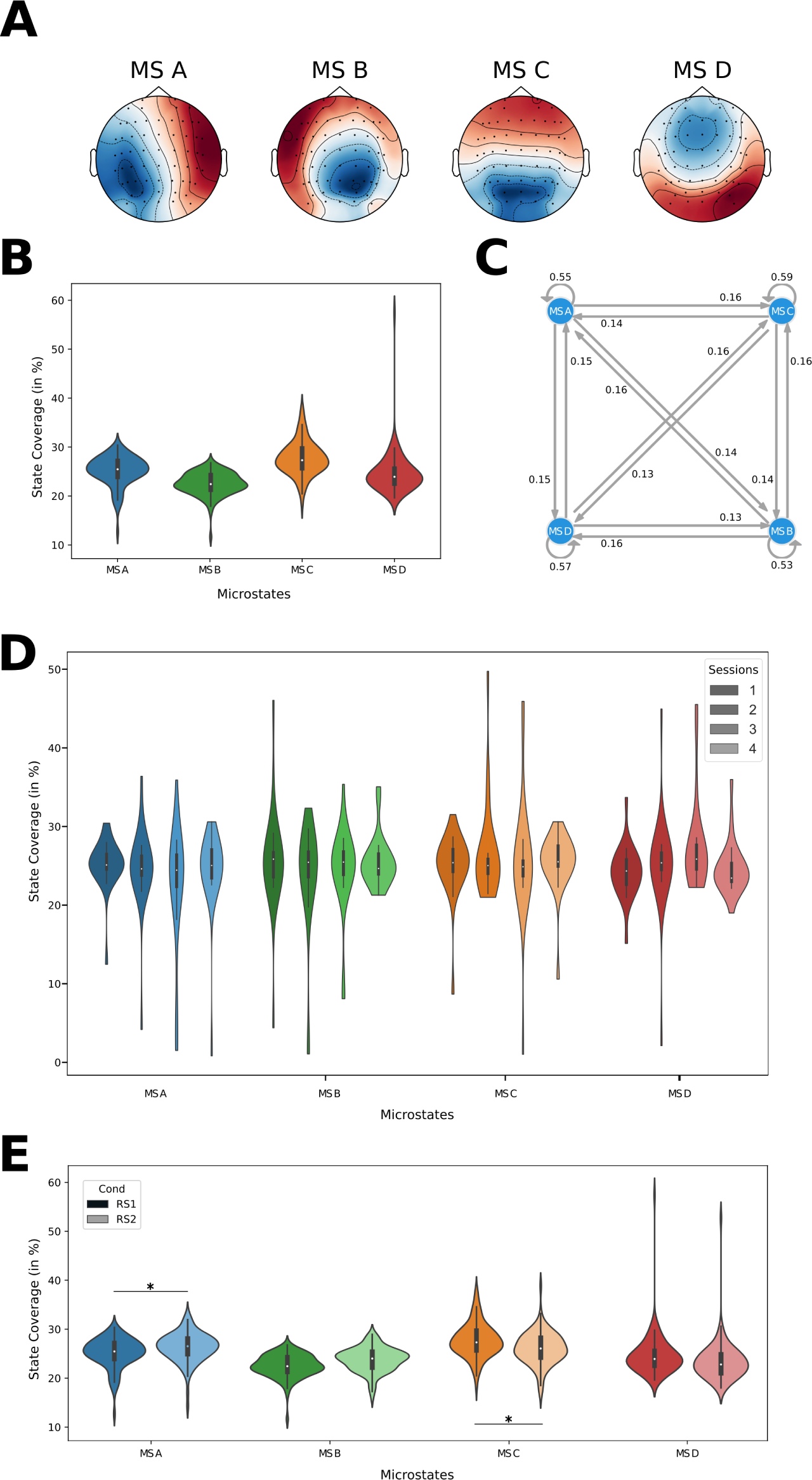
Microstates. A: Scalp microstate topographies (note: polarities are to be neglected), B: relative time the brain stays in each state, C: transitions between the four microstates displayed as a Markov diagram (A-C: all subjects, all sessions, RS1 only), D: comparison of the coverages for all sessions of RS1 across all subjects. The violins are colorcoded from dark session 1 to light session 4, E: comparison of the coverage for RS1 and RS2 across all subjects and all sessions. A darker shade represents RS1, while a lighter shade denotes RS2.

The overall picture remained unchanged when comparing the coverages of microstates between RS1 and RS2 (Fig. 3D). The frequency of each microstate was in the range of 20% to 30%. A statistical comparison also revealed no significant differences for state B (RS1: 23.78% *±* 2.86, RS2: 23.92% *±* 2.44, *p >* 0.05, *t* = *−*0.545, *df* = 95) and for state D (RS1: 24.98% *±* 4.44, RS2: 26.14%*±*3.27, *p >* 0.05, *t* = *−*1.834, df = 95). However, the other two states showed slight differences between RS1 and RS2. While state A was significantly increased in RS2 (RS1: 24.48% *±* 2.54, RS2: 26.44% *±* 2.94, *p <* 0.0001, *t* = *−*6.812, df = 95), state C was less frequent in RS2 (RS1: 26.77% *±* 3.26, RS2: 23.50% *±* 3.98, *p <* 0.0001, *t* = 5.621, df = 95).

### Source Connectivity Analysis

#### ciPLV on Source Level

After transforming the scalp-recorded EEG data to source-localized activity using dSPM implemented in MNE-Python (32), we extracted source activity from 150 parcels as defined by the Destrieux atlas (35). The proper functioning of the source localization and reasonable selection of the parcellation was ensured by using the tapping condition as ground truth for source activity (see Supplementary Material, Fig. S3). We applied a sliding window approach for the source connectivity analyses in RS1 and RS2. The mean connectivity over all subjects and sessions ranged from 0.3 to 0.6 for the different pairs of source parcels (see Supplementary Material, Fig. S4). This non-zero source-space phase-connectivity was utilized in the following to construct the networks that defined the source connectivity states of the resting brain.

#### Connectivity State Analysis

As outlined in the Materials and Methods, we set the number of clusters to *k* = 4. Additionally, we defined *k* = 8 connectivity states determined by the elbow method (Results are shown in Supplementary Material, Fig. S5).

When displaying the centroids of the clusters as circular connectivity networks and mapping them onto a 3D image of the brain, the representations resemble those of the scalp topographies of the microstate analysis (Fig. 4). The four clusters can be described as (i) a diagonal connectivity pattern of left-frontal regions (CS A), (ii) a mirrored diagonal connectivity pattern of right-frontal regions (CS B), (iii) a parietal connectivity pattern (CS C), and (iv) a weaker all-to-all connectivity pattern (CS D; max 0.1 compared to 0.2-0.35).

**Fig 4.**
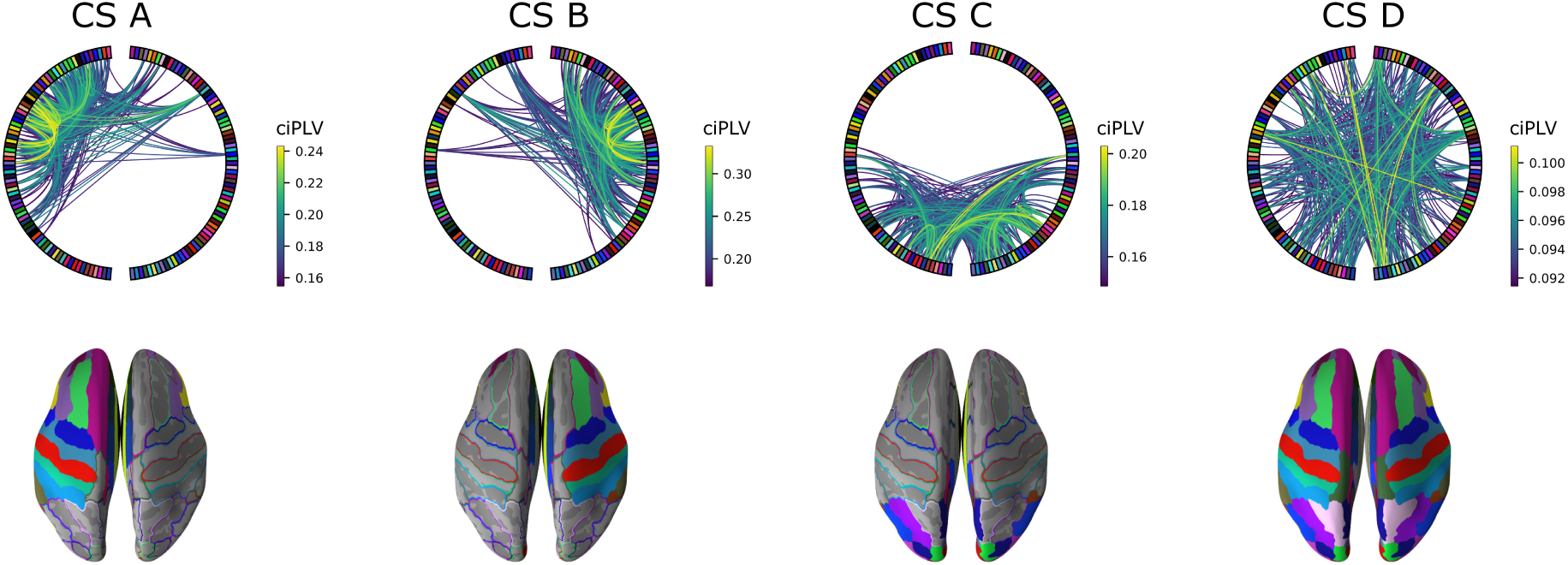
Source Connectivity States. Top: circular presentation of source connectivity (10% strongest connections shown) presenting parcels on each hemisphere on the left and right side of the circular plot, frontal parcels at the top and occipital parcels on the bottom; Bottom: presentation of the involved parcels mapped onto the brain using the same color code as in the circular plot.

In the following, we investigated the coverage of each connectivity state in the whole dataset, i.e., across all participants and measurement days (for other connectivity state metrics, please refer to the Supplementary Material, Fig. S6). For roughly 55% of the time, the resting-state network remained in connectivity state D, whose coverage was significantly increased compared to the other connectivity states (Fig. 5A and Tab. 1). Furthermore, CS B was significantly reduced compared to CS A and CS C, while CS C was significantly increased compared to CS A (Fig. 5A and Tab. 1). The states switched between CS A - CS C only about 6 *−* 10% but preferably first went back to CS D with a chance of 35 *−* 39%. This finding means that the connectivity state D served as some transition state having overall lower connectivity (Fig. 5B).

**Fig 5.**
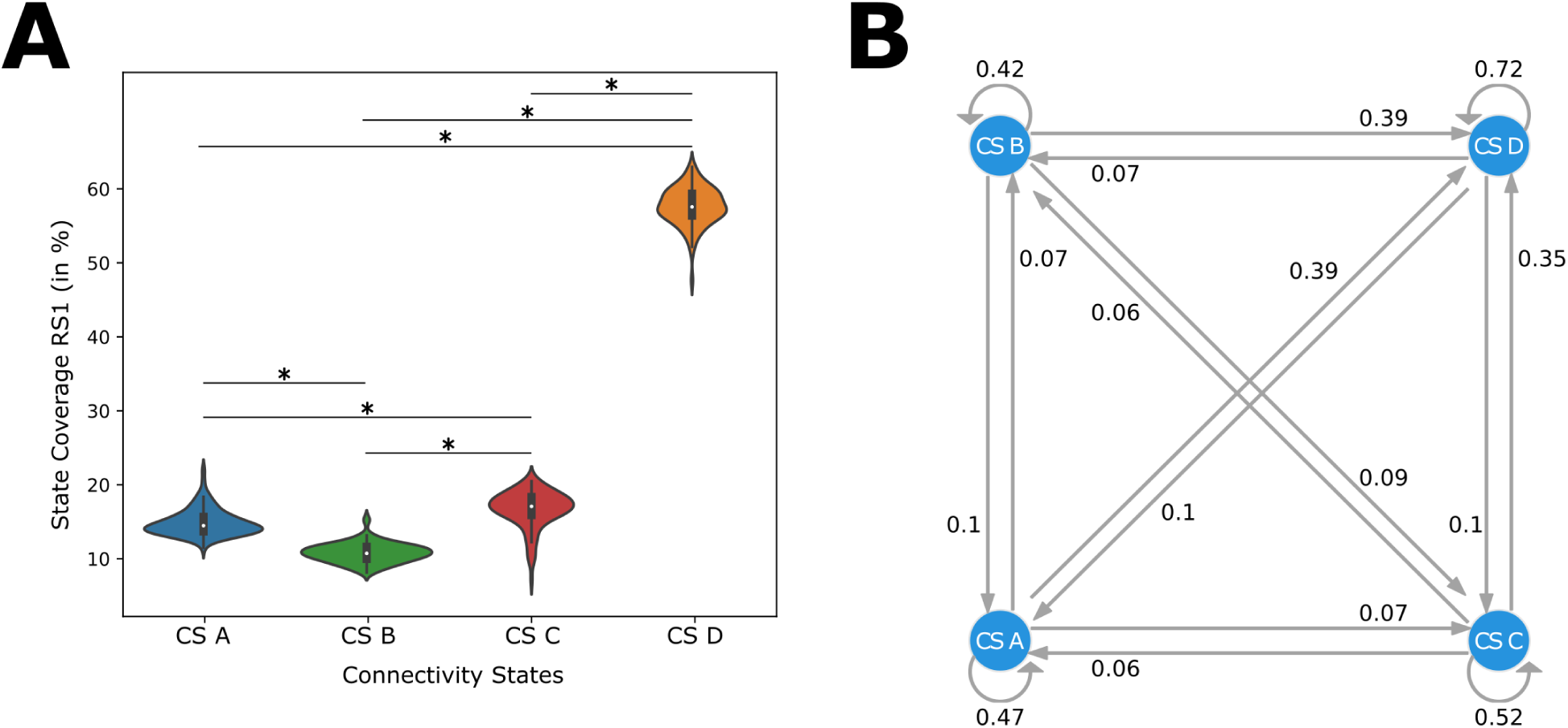
Coverages and Transitions of Source Connectivity States. A: A violinplot showing the source connectivity state coverages for CS A (blue), CS B (green), CS C (red) and CS D (orange) in RS1. All significant differences are marked with an asterisk; B: graph displaying the transitions between the connectivity states.

**Table 1.**
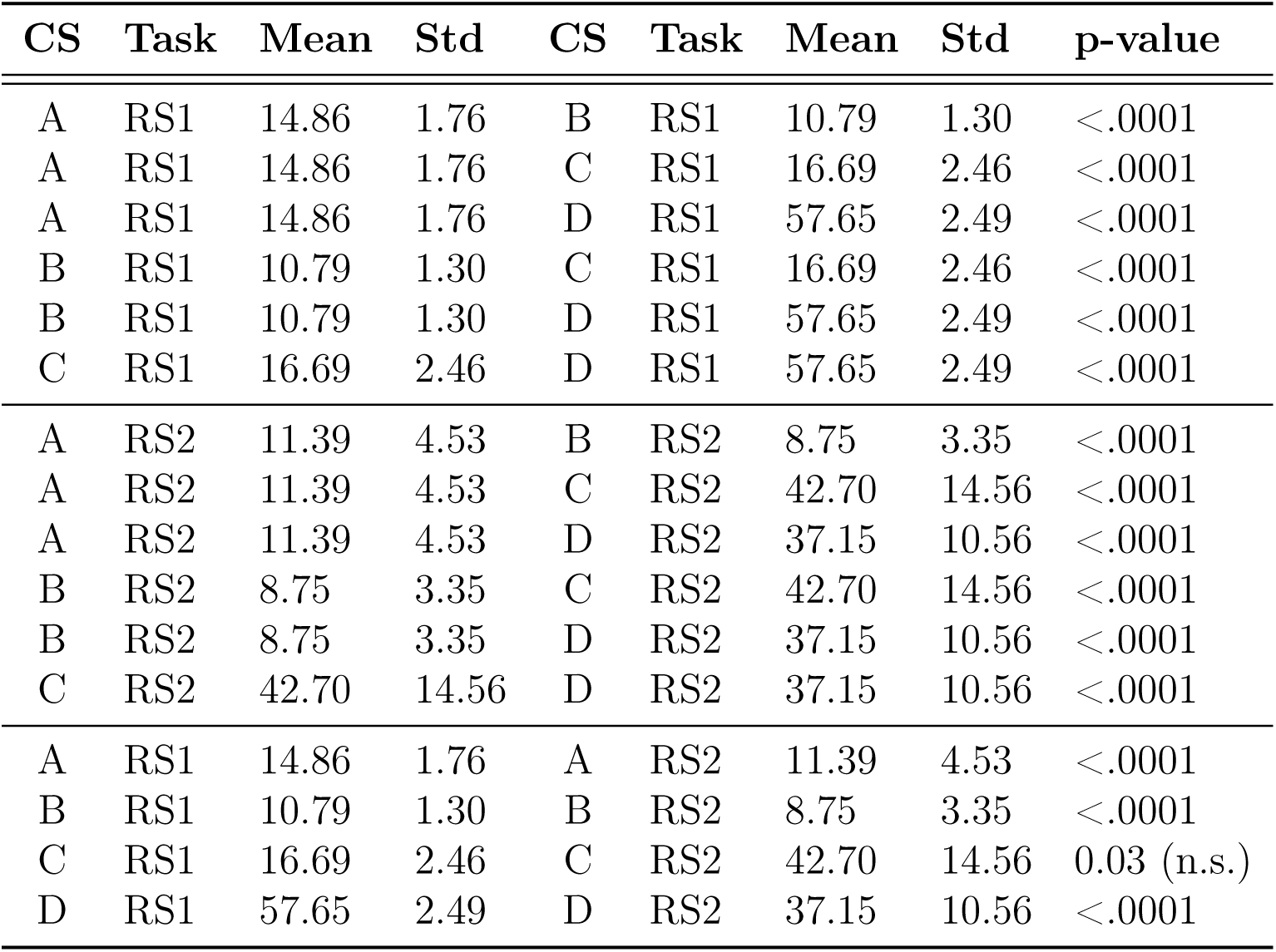
Wilcoxon signed-rank test results for the comparison of CS A, B, C and D for resting-state 1 and resting-state 2; n.s.: not significant after correction for multiple comparisons.

When separating the data for the multiple measurement days (session 1 through session 4), the same distribution between the different connectivity states could be observed (Fig. 6). Statistically, there were no significant differences (ps *>* 0.05).

**Fig 6.**
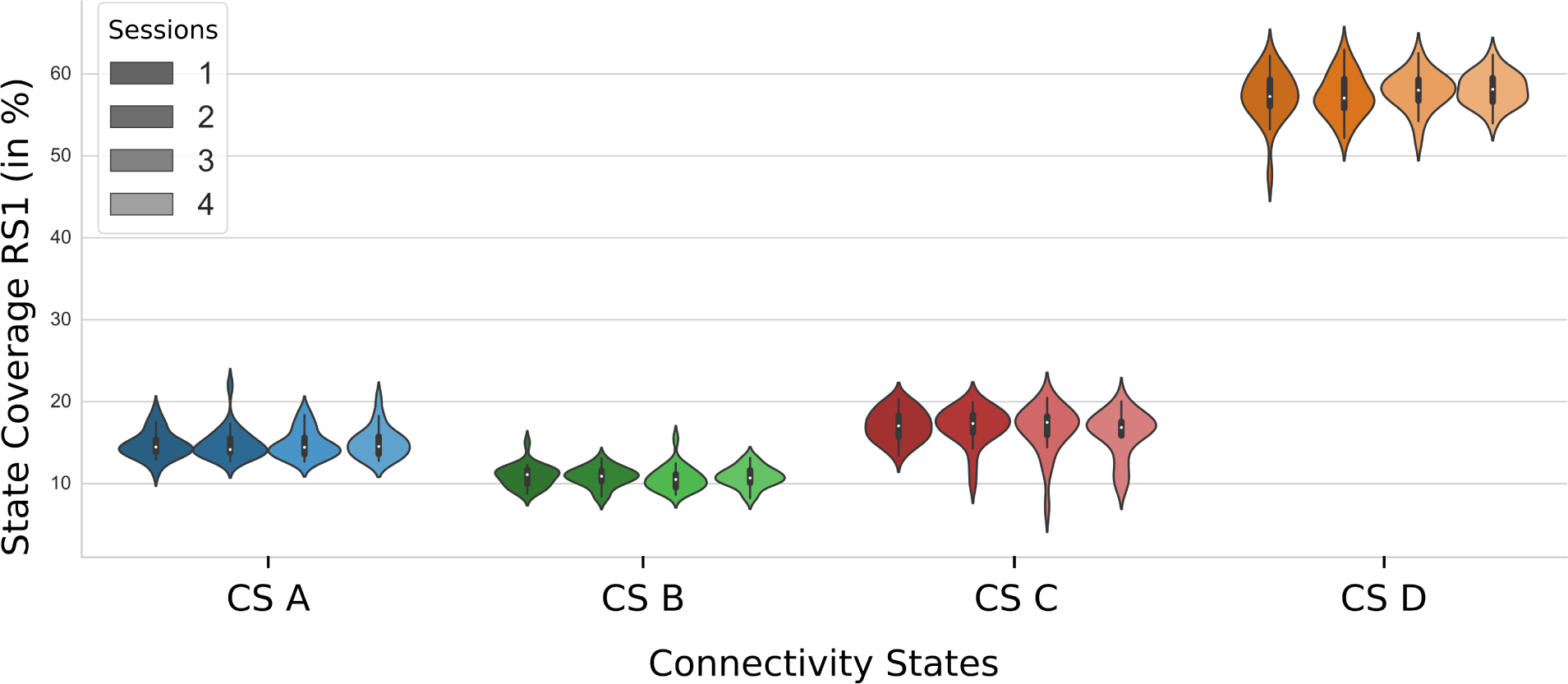
Connectivity States in RS1 for multiple measurement days. A violinplot showing the source connectivity state coverages for separate measurement days, i.e. sessions; color coding representing sessions 1 (darkest) to 4 (lightest).

Another question we wanted to answer was whether the performance of a motor task before the resting-state influences the identified connectivity states. Therefore, we compared the resting-state networks obtained from RS1 before the motor task performance to RS2 after the motor task performance. Here, CS A - CS D distributions deviated significantly between RS1 and RS2 (Fig. 7A). In contrast to the results for RS1 presented above, the coverage of CS C was significantly increased while the coverages of the states CS A, CS B and CS D were significantly reduced (Tab. 1). The significant difference between CS C and CS D in RS1 reported above vanished in RS2. The remaining significant differences, as seen in RS1, remained present in RS2 (Tab. 1).

**Fig 7.**
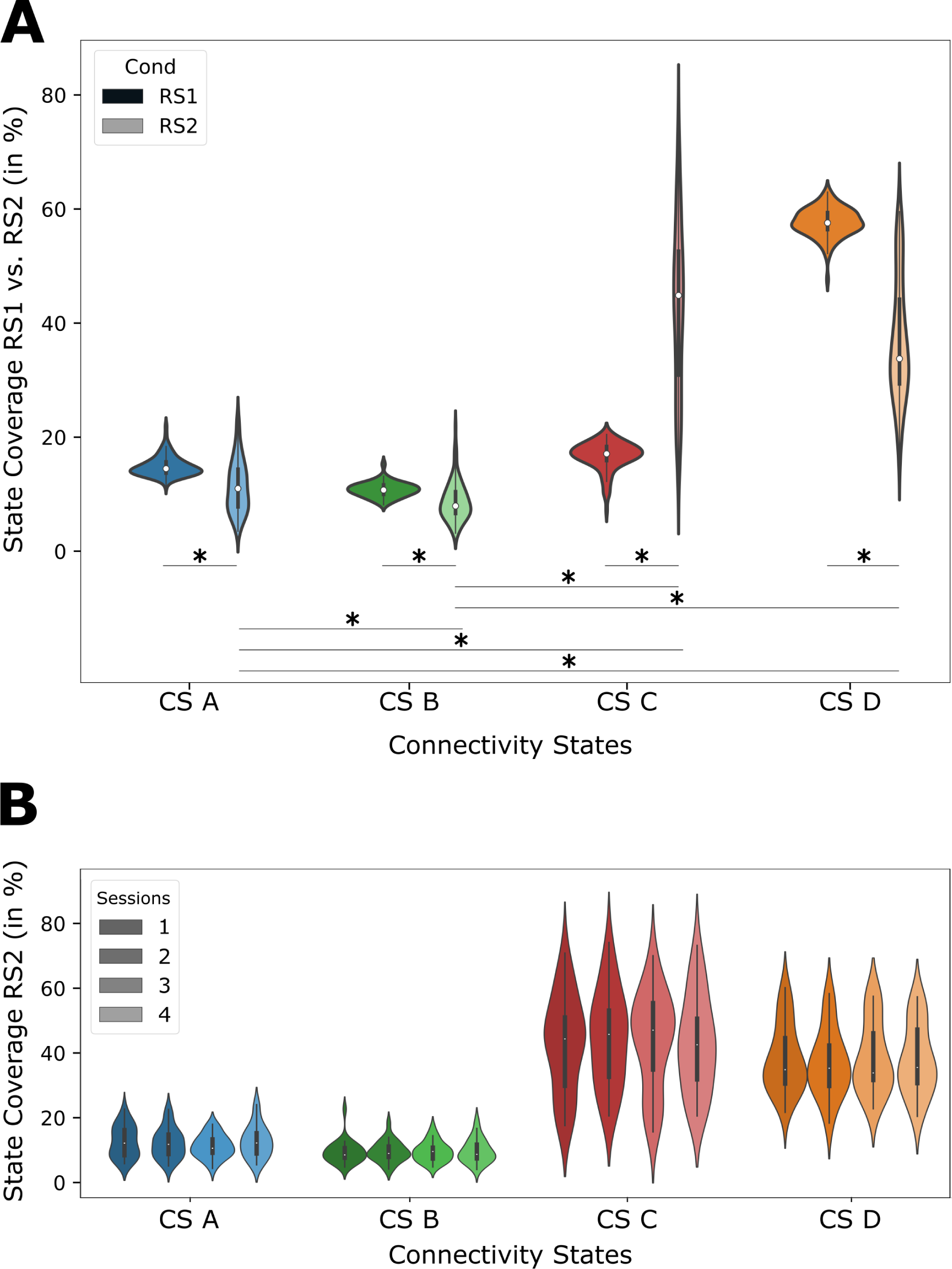
Task-dependent changes in connectivity states. A: A violinplot showing the source connectivity state coverages separately for RS1 and RS2 (all sessions); B: Stability of source connectivity state coverages in RS2 for multiple measurement sessions; color coding representing sessions 1 (darkest) to 4 (lightest).

Our results were reproducible and stable across multiple measurement days. The details are summarized in Table 2. The table presents the Intraclass Correlation Coefficient (ICC) values and corresponding *F* -values for each of the four connectivity states (CS A, CS B, CS C, CS D) in Resting State 1 (RS1, Fig. 6) and Resting State 2 (RS2, Fig. 7B). The ICC values, serving as indicators of the consistency in connectivity patterns, were calculated for each state across the 4 measurement days for both resting-states. Notably, all ICC values were found to be substantial, with RS2 exhibiting slightly higher ICC values than RS1. These findings highlight the robust reproducibility of the connectivity states across multiple measurement days and underscore the stability of our derived network representations across different resting-state sessions. The *p*-values below 0.001 further support the statistical significance of the observed reproducibility.

**Table 2.** ICC (Intraclass Correlation Coefficient) for connectivity states A-D for resting-state 1 and resting-state 2; Degrees of Freedom (*df* 1 = 23, *df* 2 = 69); all *p*-values were below 0.001.

|  | CS A |  | CS B |  | CS C |  | CS D |  |
| --- | --- | --- | --- | --- | --- | --- | --- | --- |
|  | ICC | F-Value | ICC | F-Value | ICC | F-Value | ICC | F-Value |
| RS1 | .892 | 9.22 | .753 | 4.049 | .853 | 6.793 | .842 | 6.344 |
| RS2 | .935 | 15.399 | .934 | 15.24 | .968 | 31.476 | .966 | 29.425 |

## Discussion

In this study, we conducted a source connectivity state analysis in a cohort of 24 young and healthy participants. While we performed a classical microstate analysis first, we focused not on directly comparing the scalp-level microstates and the source-level connectivity states but elucidating underlying active neural networks during resting-state using both methods. Furthermore, we wanted to establish a methodological approach that demonstrates the reproducibility and reliability of the presented method. This emphasis on methodological validation lays the ground for future applications, particularly clinical studies.

It is crucial to note that the analysis of both sensor and source space data provides valuable insight despite different numerical transformations of the same underlying data set. In particular, we gained insights into how global patterns of resting-state activity manifest at a localized level, potentially uncovering unique features and relationships that are not readily apparent from scalp-level analysis alone (45).

By establishing the connectivity state analysis and including individual structural MRIs in the source reconstruction pipeline, we anticipate this to be particularly relevant in studies of stroke patients. In particular, the ability to define individual connectivity patterns on a single-subject source space level holds promise for future investigations of stroke patients. Such applications allow for elucidating a lesion’s impact on the resting-state networks and potentially inform personalized treatment and rehabilitation strategies.

### Microstate analysis

We performed a classical microstate (MS) analysis and found a set of four MS that matched previously reported classes (e.g. (10–12)). Interestingly, we did not find any significant differences between the coverages or the transitions of the MS (Fig. 3 B-C). Furthermore, the MS were stable over multiple measurement days (Fig. 3 D).

### Connectivity state analysis

After source localization of the recordings of RS1 and RS2, the continuous resting-state data were clustered into epochs of four distinct connectivity states (CS). When comparing the CS with the MS topographically, we observed that both the MS based on scalp data and the CS based on source reconstructed connectivity networks showed similar configurations, i.e., two states with a left frontal - right parietal diagonal separation and right frontal - left parietal separation, respectively, and a state with an occipitally centered separation. The last state defined by both approaches revealed a polarity change at central electrodes in the MS, reflected by a state of all-to-all connectivity with lower connectivity strength in the CS (Fig. 4).

Given the choice of the alpha-frequency range for the analyses (see Materials and Methods), one might assume that there may be an artificially increased tendency for occipital connections as occipital alpha is dominant in resting-state EEG (46). However, our results showed the brain’s preference to stay in the intermediate all-to-all connectivity state (CS D), which we called the transition state. The transitions between the states supported this view. In only a few cases, the resting brain switched between states CS A, CS B, or CS C directly. The most frequent way to get from one state of higher connectivity to another, i.e., switch between CS A, CS B, or CS C, was to first transition to state CS D, i.e., continuously lowering overall connectivity. Thus, all states showed the highest transition probability towards connectivity state D, while the transitions back to the other states from CS D were evenly distributed. This result was stable over the multiple measurement sessions. A similar transition pattern of brain states has been reported by (47). In fMRI data analyzed with Bayesian switching dynamical systems models, which estimated the probability distribution of the system’s state based on differential equations using Bayesian statistics, they showed that states related to high cognitive load or fixation only were not suddenly shifting to one another, but first accessed states associated with an intermediate demand (47).

### Differences induced by a motor task

By including a finger-tapping task in our study, we investigated the differences in MS and CS induced by a motor task. Even though the influence on the MS coverages was significant, the change induced by the task was relatively small (MS A increased by 2%, MS C reduced by 3%). All four states were almost evenly distributed, with 25% coverages in both RS1 and RS2. In the CS, the change in the coverage of the states was much more accentuated. The coverage of CS C significantly increased by more than 25%, while all other states, especially CS D (by more than 20%), decreased significantly. The significantly reduced CS D may be related to the default mode network (48) or parts of the cognitive control network (11), which supports the view that CS D may serve as a transition state. MS D, on the other hand, is commonly assumed to be related to the dorsal attention network (11; 48). According to (49), MS D might reflect reflexive aspects of attention, as subjects underwent a lengthy motor task after which they tried to relax without focusing on the previous motor task. Assuming connectivity state C represents a similar function as microstate D, increasing this connectivity state could represent an equivalent shift of attention. Besides, the switch to a resting state after motor task performance might reflect the suppression of fronto-central activity involved in motor planning and execution (50). Furthermore, the dorsal attention network modulates the visual network (51). The frequent change of visual information presented in the motor task, which was used to inform the subjects when to start and end the finger-tapping movements, was removed in RS2. During finger-tapping, the subjects had to focus on the visual input to perform the motor task. As this attentional demand was removed in RS2, an activity change of the dorsal attention network may have resulted. We assume that the reduction of these parts of the default mode network in favor of the dorsal attention network reflects a shift from the activated motor state to a resting state and thus leads to a shift in the dorsal attention network in contrast to the pre-motor task resting-state condition.

Importantly, our data suggest that resting-state data before and after task performance should be analyzed strictly separately.

While our study predominantly focused on unraveling the impact of a motor task on the connectivity states, note that other tasks might influence the CS differently. The selection of a motor task stemmed from our interest in clinical populations with motor impairments, such as stroke patients. Nevertheless, various tasks may introduce distinct neural dynamics and connectivity patterns. As a future avenue of investigation, incorporating a range of tasks could be instrumental in achieving a more comprehensive understanding of task-induced variations in connectivity states. By examining the effects of different cognitive and motor tasks, researchers might gain deeper insights into the versatility and adaptability of brain connectivity networks, thus enriching our comprehension of their dynamic nature.

### Reproducibility

In the literature, many articles have reported a change in MS features (e.g., coverage, occurrences, transitions) due to neuropsychiatric disease (18–23). In order to extend the MS analysis to include CS of the resting brain, we aimed to show that the observed effects are stable across multiple days. We, therefore, split the recordings into four measurement days and tested the distribution of the state coverages. The results showed no significant differences within the subjects for multiple sessions. This finding suggests a reliable metric that can be applied in other populations to test for changes in CS features due to neurological disorders. For future research, e.g., in a clinical setting, this is a crucial step to provide reliable and stable results for subjects measured on different days.

### Stability of Optimal State Number Across Days

A core component of our study revolved around understanding the underlying dynamics of neural microstates. While our primary analysis focused on the 4 CS model, given its clearer interpretability, the potential variability in the optimal number of states across days for each subject remains an intriguing aspect not explored in depth in this study. The stability of the optimal number of states is a crucial metric, as it can shed light on the temporal consistency of neural network configurations in individuals. If an individual consistently exhibits a similar number of states across different days, it might suggest a more stable and intrinsic neural configuration unique to that individual. Conversely, variability in the number of states across days might indicate a more fluid neural landscape, potentially influenced by factors such as daily experiences, mood fluctuations, or cognitive demands. It’s also worth noting that while our primary focus was on the 4 CS model, our preliminary observations with the 8 CS model highlighted some markedly similar states. This finding raises the question of whether certain states might merge or bifurcate depending on the specific day or the individual’s cognitive state. Investigating this could offer insights into the hierarchical organization of these states and how they evolve. Furthermore, understanding the stability of state numbers can have broader implications. For instance, if specific individuals consistently exhibit a higher number of distinct states, it might indicate a more complex and diverse neural repertoire, which could be linked to cognitive flexibility or resilience to neural perturbations. In future research, a systematic analysis to determine the optimal number of states per-subject across multiple days would be valuable and provide a clearer picture of the temporal consistency or variability of neural state configurations.

### Limitations

While our choice of a 10% threshold for the retention of the network connections was carefully selected to strike a balance between network sparsity and representation, it is essential to acknowledge that varying threshold values could potentially lead to alterations in the network’s structural characteristics. Investigating the influence of different threshold levels on network properties represents an avenue for future research, enabling a more comprehensive understanding of the robustness and sensitivity of our derived connectivity networks.

Furthermore, it is important to mention that our sample size of 24 participants may impose constraints on the broader generalizability of our findings. Despite mitigating this limitation to some extent by obtaining measurements from each participant across four consecutive days, i.e. working with 120 data sets, the cohort size warrants caution when extrapolating the results to larger populations. As such, we recommend that forthcoming investigations consider employing larger participant cohorts to validate and expand upon the study’s outcomes, thereby enhancing the reliability and applicability of our findings in diverse contexts.

## Acknowledgments

This work was funded by the Deutsche Forschungsgemeinschaft (DFG, German Research Foundation) — Project-ID 431549029, SFB 1451, and 491111487. We thank Hannah Kirsten, Alexandra Kurganova and members of the INM-3 for their valuable assistance in data acquisition.

## Data Availability Statement

The data that supporting this study’s findings are openly available in Jülich-DATA at https://doi.org/10.26165/JUELICH-DATA/PV2LY1.

## Supplementary Material

**S1.**
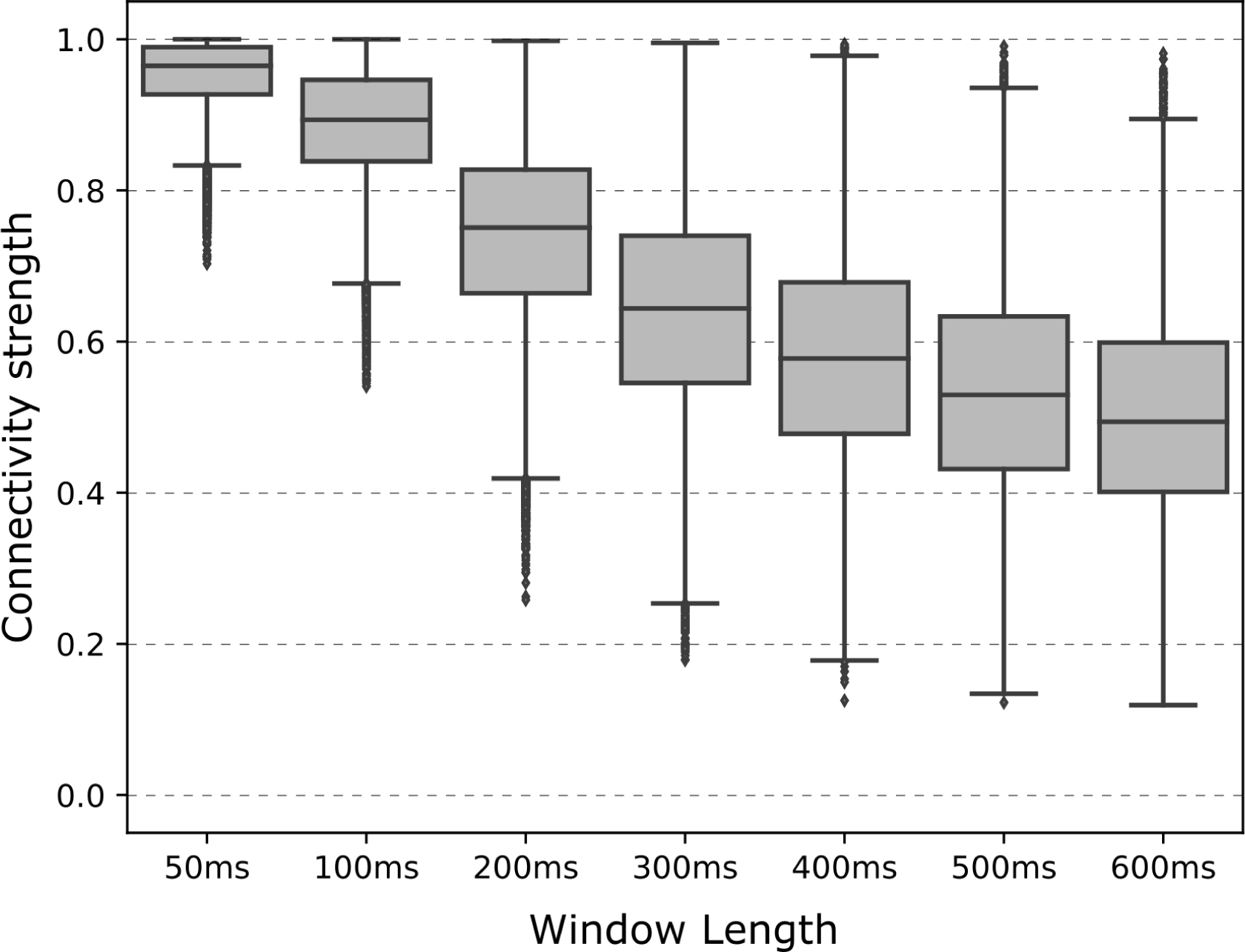
Choice of sliding window size. In optimizing our sliding window approach, selecting an appropriate window length played a crucial role in capturing the dynamic nature of the neural connectivity networks. To determine the optimal window length, a series of calculations were performed for varying window sizes, ranging from 50 ms to 600 ms, as depicted in Fig. S1. In the pursuit of aligning the window size with reported microstate coverages and leveraging the high temporal resolution of EEG data, a shorter window size initially seemed desirable. However, a notable trend emerged upon closer examination of the results obtained with the shorter window lengths. In particular, it was observed that within the alpha frequency range, almost all potential connections exhibited values approaching the maximum limit of 1. Consequently, this phenomenon led to a limited variation in the derived networks, thereby hindering the ability to discern meaningful connectivity patterns. Given these observations, a judicious compromise was necessary to balance variance and connectivity strength within the derived networks. After careful analysis, a window length of 300 ms emerged as an optimal choice. This decision was informed by the distinctive sigmoid-like trend in the mean connectivity strengths, with the 300 ms window length at this curve’s inflection point. By adopting this window length, we effectively captured dynamic variations in connectivity while maintaining a level of variance conducive to meaningful network analysis.

**S2.**
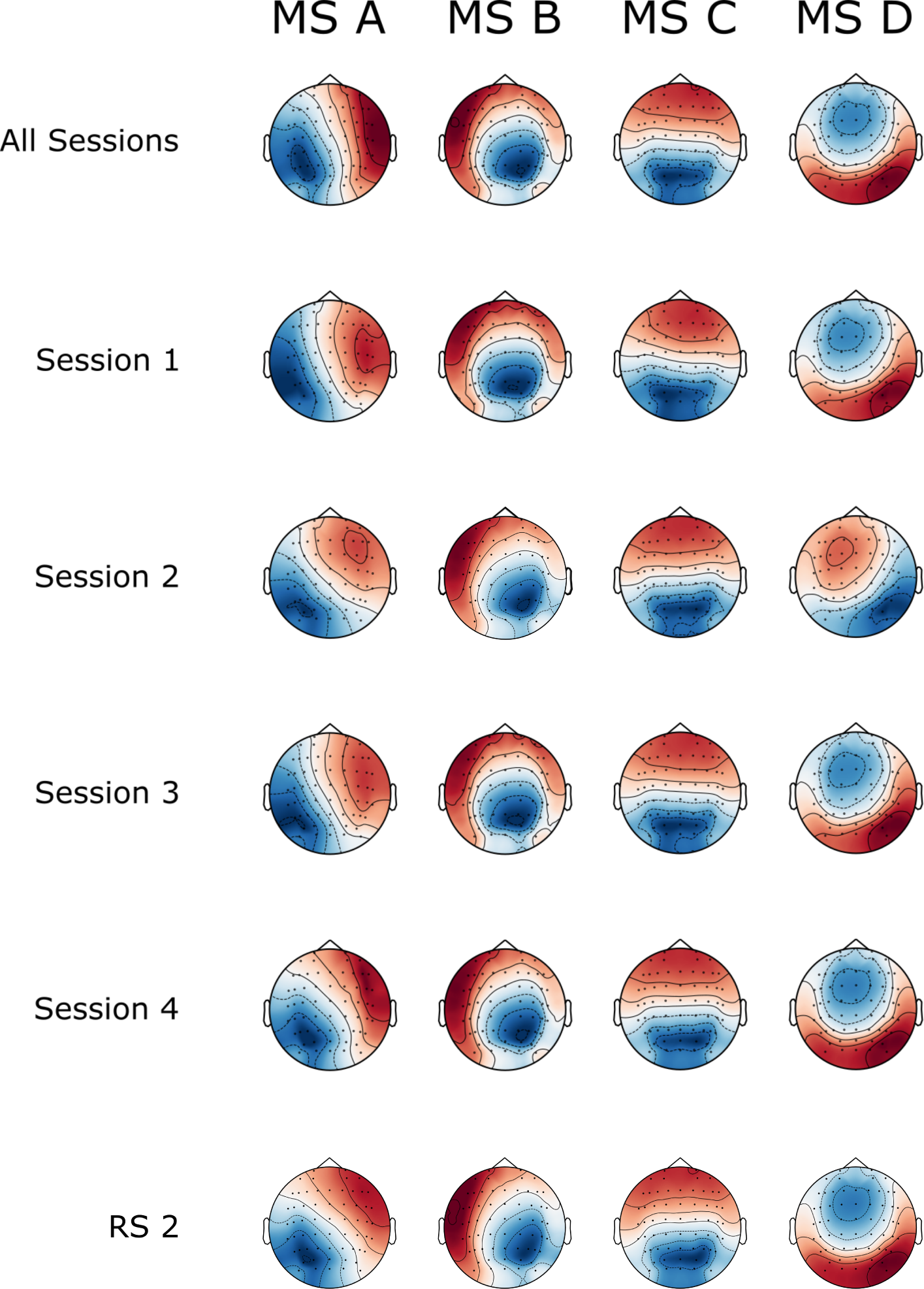
Microstate topographies remain stable over measurement sessions. In the main text, we presented the four MS topographies as defined by all four sessions combined. To check whether these states are stable across all measurement sessions, i.e., days and between RS1 and RS2, we performed the MS clustering separately for each measurement day. When comparing the MS topographies in the main text (Fig. S2, first row, RS1) with the other four MS topography configurations (second to fifth row, session 1 - 4, RS1) and RS2 (sixth row), there is a high level of similarity with only minor changes in the MS topographies. Therefore, we can assume that the MS topographies do not change significantly over the measurement days.

**S3.**
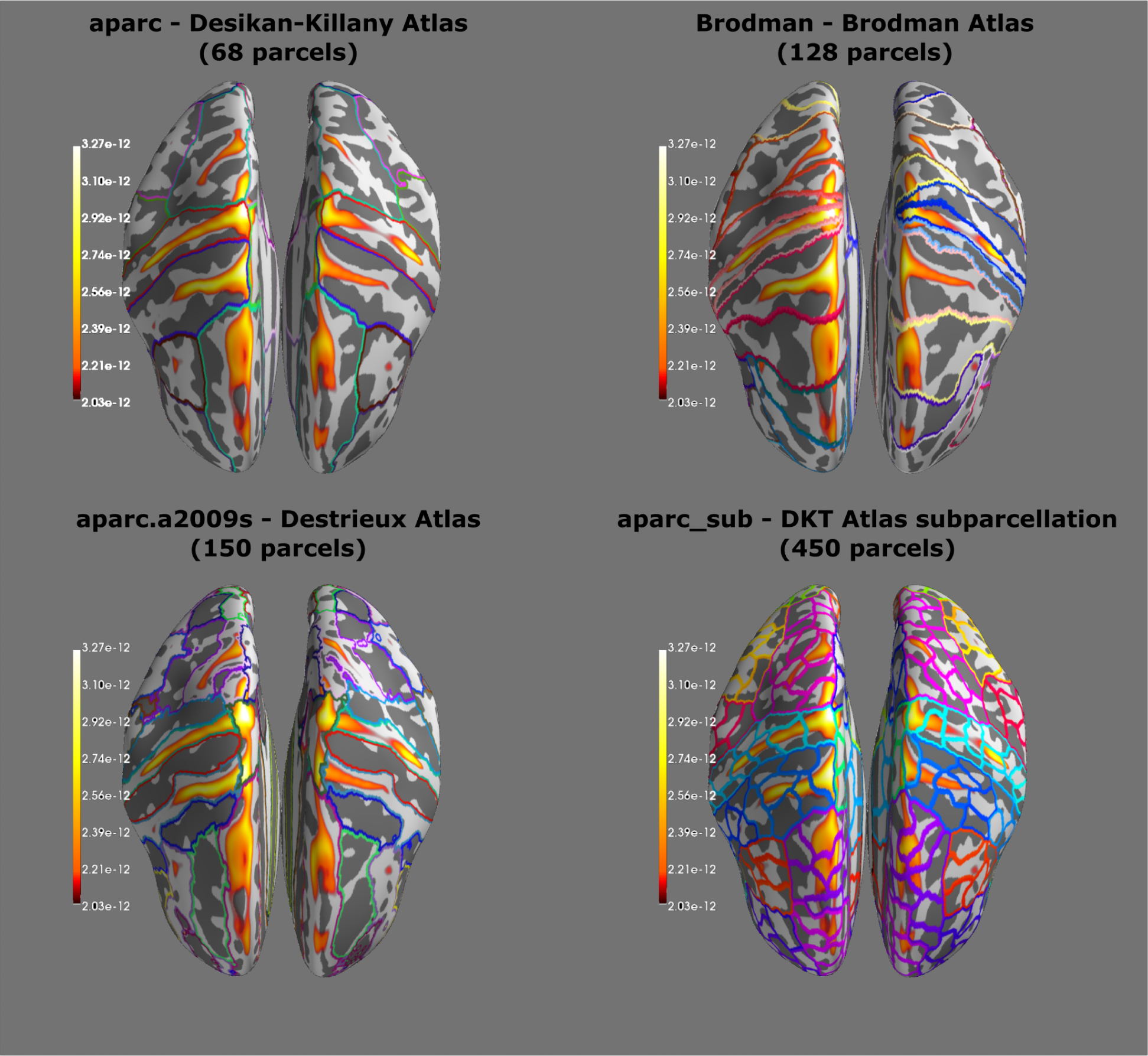
Source reconstruction - motor task. To ensure the validity of the source localization in the resting-states, we first performed a localization of the motor activity during finger movement in the "Tap" task. As shown in Fig. S3, there is an increase in activity in the area of the primary motor cortex during finger movements. Furthermore, we used this experimental condition to determine the best possible parcellation for our data. The aim was to map the activity to as fine structures as possible, which are close to the course of the source activity, but not to determine too many sources simultaneously since only 61 underlying EEG recordings were available. While the Desikan-Killany atlas and the Brodmann parcellation are coarse and not close to the observed activity, the sub-parcellation of the DKT atlas clearly defines too many sources. The Destrieux atlas is close to the observed activity and has hardly more parcellations than the Brodman atlas. Thus, we selected the Destrieux parcellation for our further investigations.

**S4.**
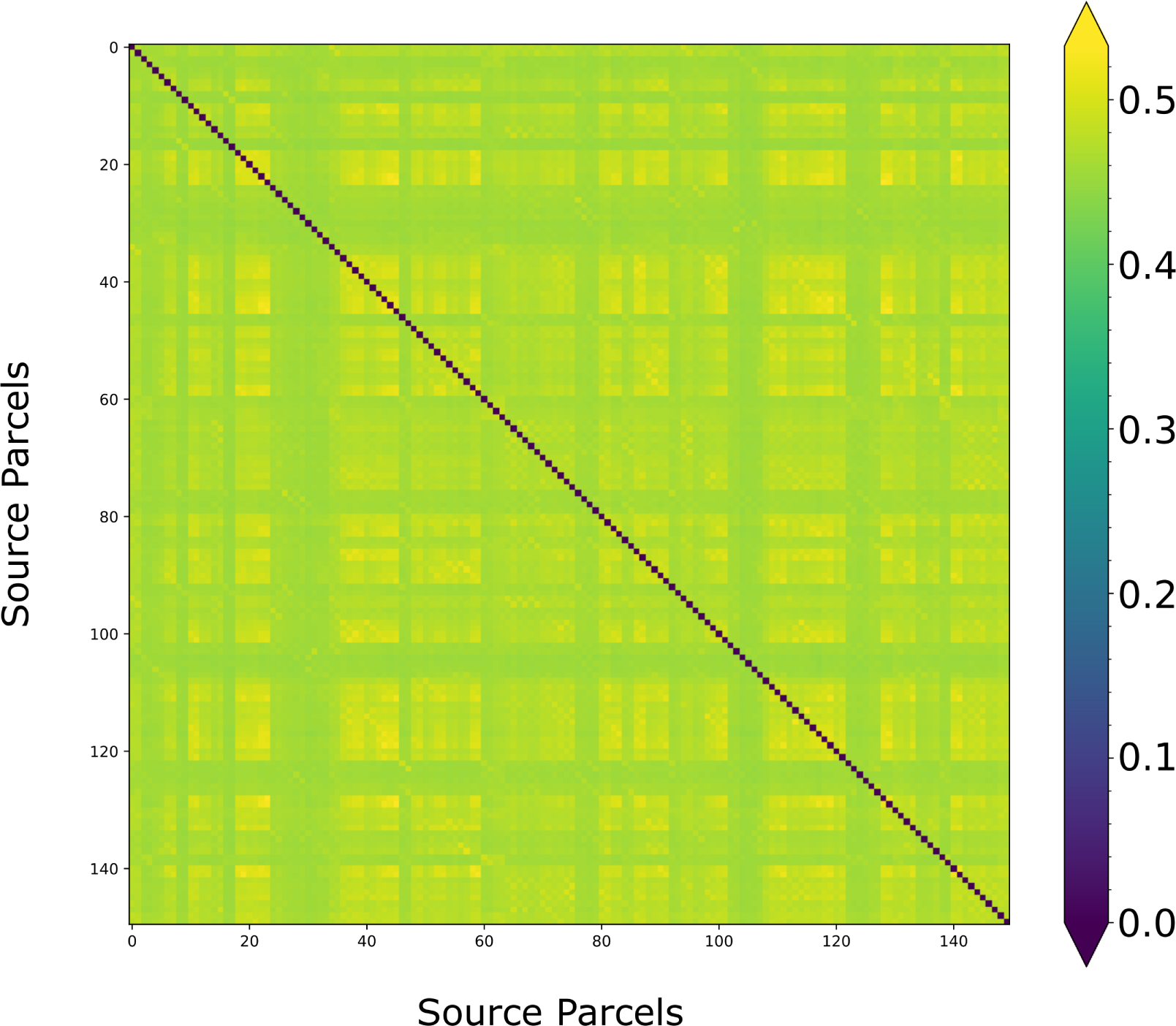
Source connectivity - resting-state. Fig. S4 provides a comprehensive exemplary view of the mean corrected imaginary phase-locking value (ciPLV) across all subjects, sessions, and time windows for each pair of source parcels under investigation. The purpose of this figure is to demonstrate the existence of varying connectivity strengths throughout the entire network, indicating that the connections between these parcels are not zero. This observation is crucial as it signifies the presence of meaningful connectivity within the network. By illustrating the non-zero connectivity, this figure supports the notion that the networks formed based on these connections carry significance and are not merely random or spurious associations.

**S5.**
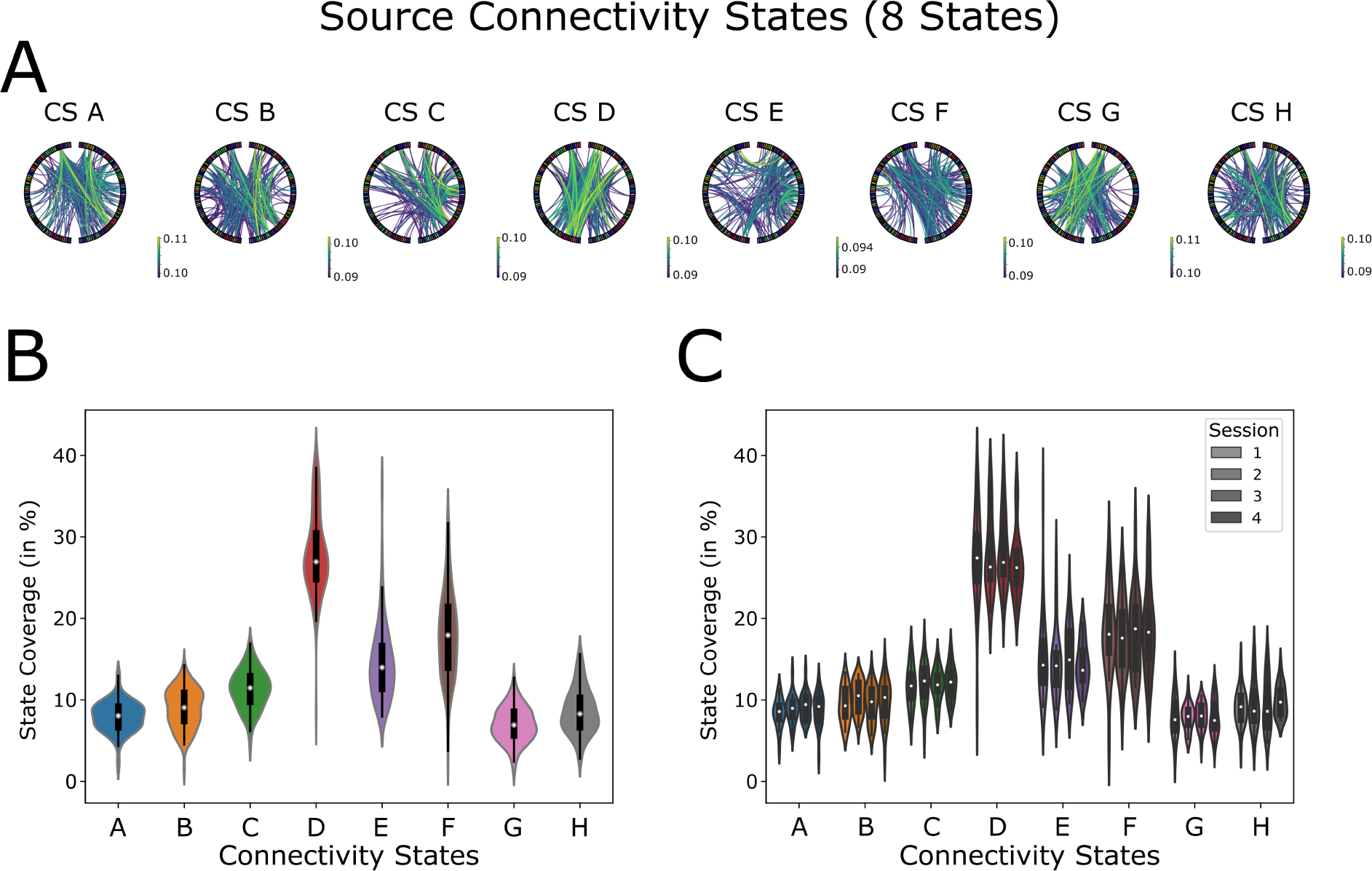
Connectivity state analysis for 8 states. Besides the analysis defining 4 connectivity states, we clustered the data into 8 8 connectivity states (CS). The results are presented in Fig. S5. We generally observed similar phenomena as in the results presented in the main text. We found 5 states (CS A, B, C, G, and H) that occurred roughly 10% of the time, while three other states showed an increased duration of 15% (CS E), 20% (CS F), and 28% (CS D). The same behavior could be observed when dividing the data into measurement sessions. When examining the patterns of the CS, it is noticeable that there are remarkable similarities to the phenomena described in the main text (e.g., CS C, E, and G). Besides the patterns already described, 4 other patterns appeared, which in some cases (such as CS A and CS C) were in part very similar to each other, and thus, one should be careful in arguing that they represent new/independent states.

**S6.**
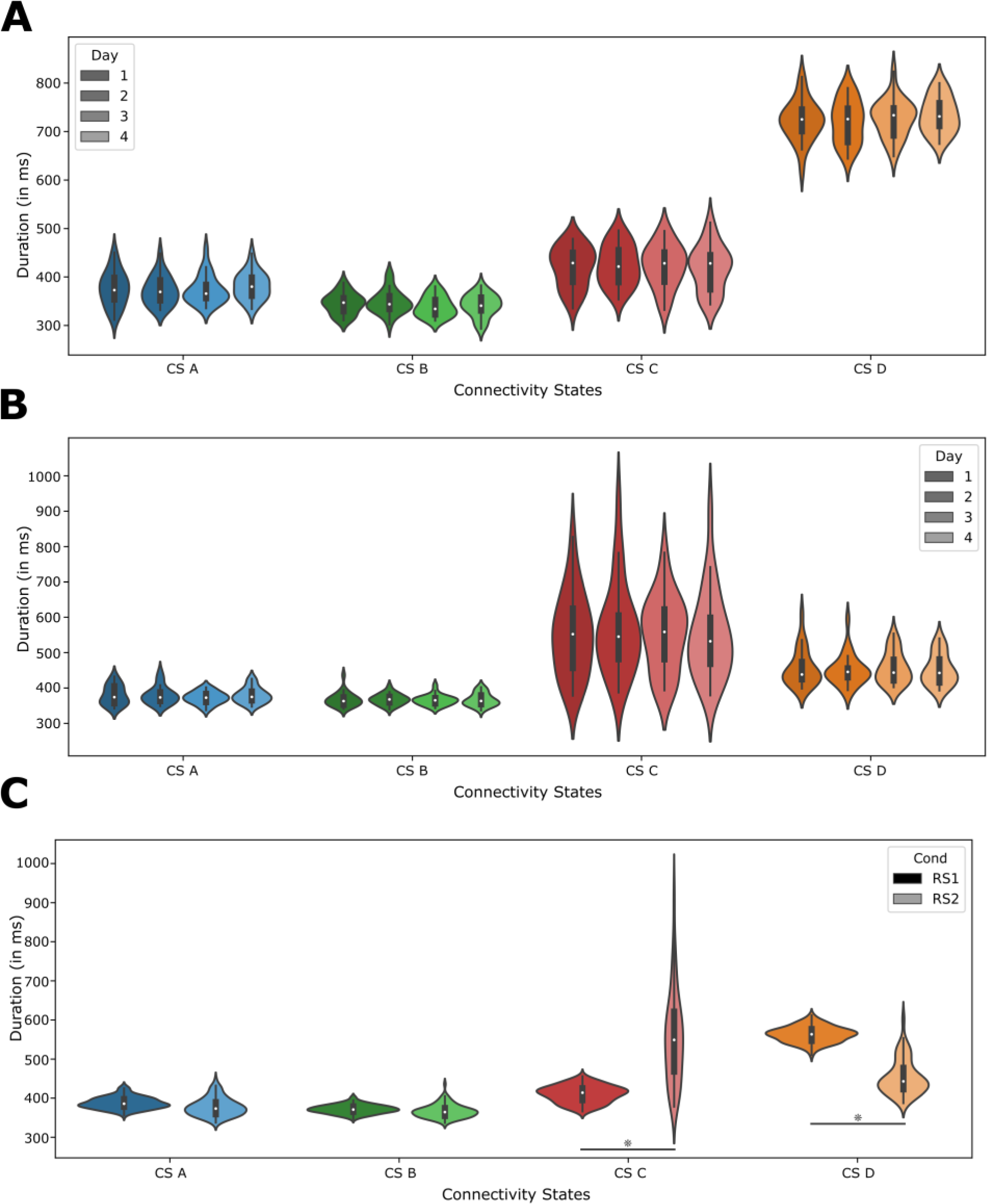
Connectivity state metrics. Additionally to our analysis of CS coverages, we explored CS frequency, contribution and duration. We observed that frequency and contribution share characteristics akin to CS duration. For this reason, we show the results on the connectivity state duration only, which is defined as the average duration for which the network persistently remains in a single connectivity state, see Fig. S6. Our observations revealed consistent patterns across all four measurement days (Fig. S6 A and B). For both RS1 and RS2, this metric was comparable to the CS coverage. Notably, in RS1, the CS D remained the dominant state in duration, mirroring its prevalence in total coverage (Fig. S6 A and C). In RS2, though the coverage indicated considerable variance for CS C, its average duration was extended, whereas the duration for CS D was diminished (Fig. S6 C). The overall consistency of this metric reaffirms the reproducibility of our methodological approach.

